# Estimating heritability explained by local ancestry and evaluating stratification bias in admixture mapping from summary statistics

**DOI:** 10.1101/2023.04.10.536252

**Authors:** Tsz Fung Chan, Xinyue Rui, David V. Conti, Myriam Fornage, Mariaelisa Graff, Jeffrey Haessler, Christopher Haiman, Heather M. Highland, Su Yon Jung, Eimear Kenny, Charles Kooperberg, Loic Le Marchland, Kari E. North, Ran Tao, Genevieve Wojcik, Christopher R. Gignoux, PAGE Consortium, Charleston W. K. Chiang, Nicholas Mancuso

## Abstract

The heritability explained by local ancestry markers in an admixed population 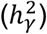 provides crucial insight into the genetic architecture of a complex disease or trait. Estimation of 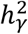 can be susceptible to biases due to population structure in ancestral populations. Here, we present a novel approach, Heritability estimation from Admixture Mapping Summary STAtistics (HAMSTA), which uses summary statistics from admixture mapping to infer heritability explained by local ancestry while adjusting for biases due to ancestral stratification. Through extensive simulations, we demonstrate that HAMSTA 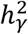 estimates are approximately unbiased and are robust to ancestral stratification compared to existing approaches. In the presence of ancestral stratification, we show a HAMSTA-derived sampling scheme provides a calibrated family-wise error rate (FWER) of ∼5% for admixture mapping, unlike existing FWER estimation approaches. We apply HAMSTA to 20 quantitative phenotypes of up to 15,988 self-reported African American individuals in the Population Architecture using Genomics and Epidemiology (PAGE) study. We observe 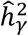 in the 20 phenotypes range from 0.0025 to 0.033 (mean 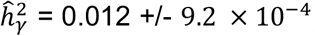), which translates to 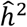 ranging from 0.062 to 0.85 (mean 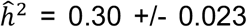). Across these phenotypes we find little evidence of inflation due to ancestral population stratification in current admixture mapping studies (mean inflation factor of 0.99 +/-0.001). Overall, HAMSTA provides a fast and powerful approach to estimate genome-wide heritability and evaluate biases in test statistics of admixture mapping studies.

## Introduction

Admixture mapping (AM) aims to identify genomic regions associated with a disease or quantitative trait in recently admixed populations^1–7^ by leveraging the differences in allele frequencies between local ancestries^8^. AM provides a powerful approach to complement genome-wide association studies (GWAS) in admixed populations due to local ancestry information better tagging uncommon or poorly imputed causal variants^5^ and spanning larger genomic regions, thus reducing the multiple testing burden^9^, enabling discoveries with relatively smaller sample sizes ^3,10^. Similarly, recent work^11^ demonstrated that local ancestry information, which is summarized by heritability explained by local ancestry 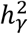, can be leveraged to estimate narrow-sense heritability *h*^2^ in admixed populations, unlike the genotype-based lower bounds (i.e. 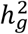). Multiple works have shown that population structure can bias association tests and estimates of 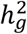 ^12,13^. However, it is less understood how similar demographic phenomena bias AM and 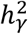 inference in admixed populations.

Admixed populations are typically modeled as a mixture of multiple continental ancestries (e.g., African, European, or Native American) with finer-scale structure within ancestral populations left unmodeled. Nevertheless, human populations are often structured across both space and time. For example, European ancestry individuals can be modeled as a mixture of at least three ancient populations^14^, and Native American ancestry components found in Latinos can also be derived across multiple subpopulations spread across Latin America^15^. This unmodeled fine-scale structure could lead to potential biases in downstream association testing. Indeed, this phenomenon has been demonstrated in European populations ^16,17^, and could similarly impact inference in admixed populations when it is not fully accounted for ^18^. When estimating *h*_*g*_^2^ using SNP data of large sample size, a robust approach to population stratification is to estimate *h*^2^ and test statistic inflation simultaneously^19^. Examples of this approach include linkage disequilibrium score regression (LDSC)^13^ and cov-LDSC^12^. While these methods are designed for SNP data, it remains unclear how applicable they are on estimating 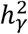 using summary statistics from admixture mapping studies.

In this study we propose HAMSTA (Heritability estimation from Admixture Mapping Summary STAtistics), a novel likelihood-based approach to infer 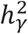 from admixture mapping summary statistics. To achieve robust and efficient computation, HAMSTA transforms the correlated test statistics using a truncated singular value decomposition (tSVD) and performs maximum-likelihood inference while accounting for residual inflation due to stratification within ancestral populations. We perform extensive simulations and demonstrate that HAMSTA provides approximately unbiased estimates of 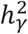 and outperforms existing approaches to detect evidence of stratification bias. We demonstrate estimates from HAMSTA can be leveraged to efficiently compute well-calibrated family-wise error rates for admixture mapping, particularly in presence of ancestral stratification which previous approaches do not consider ^20^. Next, we apply HAMSTA to admixture mapping summary statistics for 20 traits from 15,988 self-identified African American individuals in the Population Architecture using Genomics and Epidemiology (PAGE) study ^21^. We find the *h*^2^ estimates of 0.85 (0.085) and 0.42 (0.086) for height and BMI respectively. Compared with LDSC on admixture mapping summary statistics, HAMSTA offers more precise estimates for 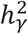 and better quantifies the inflation in the test statistics due to unknown confounding biases. Overall, we demonstrate that HAMSTA provides a fast and powerful way to estimate genome-wide heritability that controls biases using summary statistics from admixture mapping studies.

## Materials and Methods

### Model for complex trait and ancestral stratification

We consider a two-way admixed population, with ancestral populations pop1 and pop2, which is recently structured into pop2a and pop2b (**Supplementary Figure 1**). This demographic model mimics the African and European admixture in African American and the finer-scale structure in their ancestral European population. We let *γ, δ* and − *δ* denote the population mean phenotype values of pop1, pop2a and pop2b. We denote *A*_*i,k*_ as the centered and standardized local ancestry calls for individual *i* at marker *k*, such that *E*[*A*_*i,k*_] = 0and *Var*[*A*_*i,k*_] = 1. We denote indexing over *N* individuals at the *k*th marker as *A*_*k*_ and index over *M* markers for the *i*th individual as *A*_*i*_. We define the phenotype *y*_*i*_ of an admixed individual *i* as,

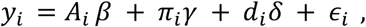

where *β* is the *M* × 1 vector of local ancestry effects, *π*_*i*_ is defined as the global ancestry proportion due to pop1, 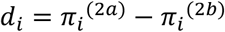 is the difference between the global ancestry proportions of pop2a and pop2b, and *ϵ*_*i*_ ∼ *N*(0, *σ*^2^_*ϵ*_) is residual environmental noise. Furthermore, we assume that 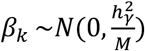, where 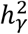 is defined as the heritability explained by local ancestry ^11^. Lastly, we define 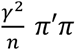 as the phenotypic variance explained (PVE) by global ancestry, and 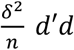 as PVE by ancestral stratification.

### Test statistics for admixture mapping

We model the marginal association statistics from an admixture mapping study where only global ancestry proportions *π*_*i*_ (and not *d*_*i*_) are known beforehand. If the stratification term is not adjusted, the test statistics for marker *k* will be *Z*_*k*_ = *s*_*R*_^−1^(*A*_*k*_^′^*PA*_*k*_)^−1/2^(*A*_*k*_^′^*Py*), where *s*_*R*_^2^is the residual variance after the global ancestry *π* is projected out by matrix *P* = *I* − *π*(*π*^′^*π*)^−1^*π*^′^. Extending this to all *M* markers we have, *Z* = *s*_*R*_^−1^*D*^−1/2^(*A*^′^ *Py*), where *D* is the diagonal elements of *A*^′^*PA*. Given this and distributional assumptions regarding *y*, we can derive the expectation and covariance of *Z* as,

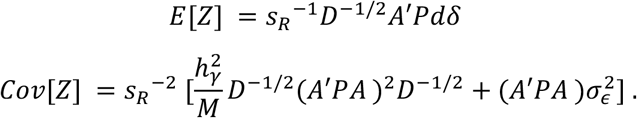

The *D*^−1/2^(*A*^′^*PA*)*D*^−1/2^ is local ancestry disequilibrium (LAD) matrix analogous to the LD matrix and *D*^−1^(*A*^′^*PA*)^2^*D*^−1^ is the LAD score matrix in which element (*j, k*) is approximately the dot product of correlation vectors of two markers *j* and *k*. When sample size N is large, the test statistics *Z* are well-approximated by a multivariate normal distribution. The mean reflects the bias due to correlation between local ancestry and ancestral stratification conditional on the global ancestry. In the covariance, the first term is related to the heritability explained by local ancestry and LAD score matrix. The second term in the covariance is related to LAD matrix and nongenetic effects. In the null scenario, where 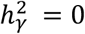, *δ* = 0, the distribution of *Z* has means of zeros and covariances simply equal to the LAD matrix.

We let the singular value decomposition (SVD) of *A*^′^*P* = *USV*^′^, *A*^′^*PA* = *US*^2^*U*^′^ and (*A*^′^*PA*)^2^ = *US*^4^*U*^′^. We define rotation *Z*^*^ = *S*^−1^*s*_*R*_*U*^′^*D*^1/2^*Z*, which follows 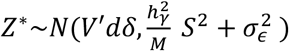, where the components are independent. We then assume *V*^′^*dδ* to be random and follow a normal distribution *N*(0, *C*^*^) such that 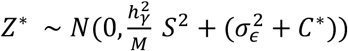. The parameters 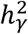 and “intercept” 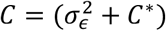 are the parameters to be inferred. To allow heterogeneous *C* across *Z*^*^, we allow *C* to be different every 500 elements, i.e., *C* = (*c*_1_ …_×500_, *c*_2_ …_×500_, …). Test statistics from different chromosomes are rotated separately and do not share elements in *C*.

### Inferring 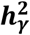 and biases using HAMSTA

Parameters 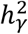 and *C* were log-transformed to ensure positivity during optimization. First, we test for ancestral stratification using a likelihood ratio test between models with multiple intercepts and single intercepts in which *C* is a scalar shared by all elements in *Z*^*^. If the test is significant with *p* < 0.05, we determine the maximum likelihood estimates 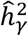 and *Ĉ* under the multiple intercept model. Otherwise, we find 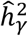 and *Ĉ* under the single intercept model. To test for the significance of 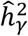, we use a likelihood ratio test that test the hypothesis 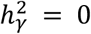. The standard errors of the estimates were determined using the jackknife method over 10 blocks.

### Estimating *h*^2^ from 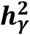

Previous work^11^ demonstrated a relationship between narrow-sense heritability *h*^2^ and 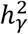 as 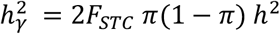. The *F* is defined as the average genetic distance between the ancestral populations at causal loci. At each site, the genetic distance is computed as 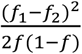, where *f*_1_, *f*_2_ and *f* are the allele frequency in the ancestral populations and the admixed population. We provided *h*^2^ estimates based on 1) *F*_*STC*_ = 0.1692 reported in the original study^11^, which was estimated from HapMap 3 dataset and 2) *F*_*STC*_ = 0.1152 estimated in this study using a subset of African and European descent from the 1000 Genome and HGDP subset in gnomAD v3.1 ^22^, assuming common variants explain 90% of *h*^2^. The average admixture proportion *π* was observed to be 78% African ancestry.

### Simulation design

To validate and assess performance of HAMSTA we performed simulations using realistic demographic scenarios. Specifically, we simulated ancestral populations *pop1* and *pop2* mirroring African and European populations in the Out-of-Africa demography model ^23^. We additionally introduced structure into *pop2* by subdividing it into two subpopulations (denoted by *pop2a* and *pop2b* below, **Supplementary Figure 1**). We set *pop2a* and *pop2b* to have diverged 200 generations ago with a migration rate = 10^−3^. These parameters were selected to result in a genetic differentiation similar to that within European populations (*F*_*ST*_ ≈ 0.003) estimated from the HGDP and 1000 Genome subsets in gnomAD ^22^. We simulated this demography for a 250Mb region with a uniform recombination rate of 10^−8^ per bp using msprime ^24^. Using the true genealogies from simulations, we extracted the true local ancestry of each individual by tracing their lineage to each ancestral population (pop1, pop2a or pop2b). Global ancestries were computed from local ancestry information by computing the total proportion of the 250Mb region that is inherited from an ancestral population. We sampled 50,000 admixed individuals and 20,000 local ancestry markers according to the demography mode.

Next, we simulated phenotypes according to our phenotype model *y* = *A β* + *πα* + *dδ* + *ϵ*. Given *α* sparsity, we drew the effect of a local ancestry marker *β*_*k*_ from 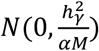 with probability *α* and *ϵ* from 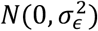. Then we set the true 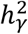, PVE by global ancestry, PVE by ancestral stratification, and 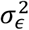 by varying the values of *γ* and *δ*. Finally, test statistics were computed using linear regression adjusting for *π* using PLINK 2.0 ^25^.

### Estimate 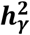 with other approaches

To compare HAMSTA with existing methods in 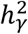 estimation, we applied BOLT-REML^26^, GCTA ^27^ and LD score regression (LDSC)^13^ to the simulated and real-world data. In GCTA, the same set of covariates included in the admixture mapping were used in 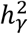 estimation. Following previous studies, we compute the genetic relatedness matrix using local ancestry in place of genotypes ^11^. In LDSC, we define the “local ancestry linkage disequilibrium” (LAD) score for marker *i* as 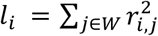 with *W* being the set of markers in a given window size. Window sizes of 1-cM and 20-cM were used. The LAD scores were used as the regressors and weights in LDSC.

### Significance threshold estimation

Specifically, to determine the significance threshold for a given admixture mapping study, we randomly generated test statistics 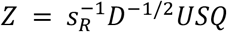, where *Q* is a vector of random variable sampled from 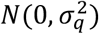. We set 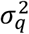 to be the maximum intercept if the test for multiple intercepts is significant, and 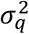 to be the inferred intercept if the test is not significant. We repeated the sampling procedure 2,000 times to determine the critical value as the 95% percentile of *max*(*Z*^2^). The significance threshold was determined as the tail probability of a chi-square distribution (degree of freedom = 1) at the critical value. To determine the threshold for multiple chromosomes, we estimate the threshold for each chromosome separately and then combine the thresholds by summing up the effective testing burden, i.e., 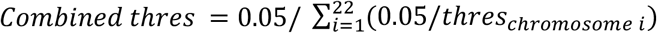. For comparison, we also estimated the significance threshold using STEAM^20^, which sampled from *Z* = *MVN*(0, ∑), where ∑ is a local ancestry correlation matrix based on genetic distance and admixture parameters. Family-wise error rates (FWER) were computed as the percentage of times at least one significant signal is identified out of 500 null simulations.

### Local ancestry inference and genome-wide mapping for admixed individuals in PAGE cohort

We obtained phenotypes and genotyping data measured on Multi-Ethnic Genotyping Array (MEGA) from the PAGE study ^21^. The complete dataset included 17,299 participants who self-identified as African American. Our analysis included 20 quantitative phenotypes: Body mass index (BMI), height, waist-to-hip ratio, diastolic blood pressure, systolic blood pressure, PR interval, QRS interval, QT interval, fasting glucose, fasting insulin, C-reactive protein, mean corpuscular hemoglobin concentration, platelet count, estimated glomerular filtration rate, cigarettes per day, coffee cups per day, high-density lipoprotein (HDL), low-density lipoprotein (LDL), triglycerides, and total cholesterol. Filters and transformations were applied, and covariates were selected according to the original PAGE analysis within the African American subset ^21^.

To infer the local ancestry, a subset of African and European genomes from the 1000 Genome and HGDP subset in gnomAD were used as reference individuals ^22^. After filtering out SNPs with missingness > 10%, lifting over and merging, 516,731 SNPs were used in the local ancestry inference, resulting in 101,292 local ancestry markers. The genotypes of PAGE and reference individuals were re-phased together using EAGLE ^28^, and the ancestry probabilities were inferred as the local ancestry of the haplotype in a region using RFMIX2 ^29^. The global ancestry of an individual was computed by taking the average of all predicted local ancestries. We analyzed up to 15,988 individuals who have >5% of one of the inferred ancestries and have no missing values in the covariates in the 20 quantitative phenotypes. Admixture mapping was performed using linear regression adjusting for the study center, inferred global ancestry, and phenotype-specific covariates used in PAGE. The average estimate of 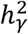 across phenotypes was calculated by weighting the estimate of each phenotype by the inverse of the squared standard error. The run time was measured on a machine with an Intel Xeon 4116 processor and 48GB memory.

## Results

### HAMSTA provides unbiased estimates of 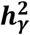 under ancestral stratification

To evaluate the accuracy of 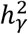 estimates under various scenarios, we performed simulation studies using local ancestry data simulated under a population demographic model that mirrors African American admixture history with an addition of recent population structure in one of the ancestral populations (see **Methods**). Briefly, we simulated phenotypes absent stratification effects where we varied 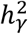 from 0 to 0.05 (corresponding to *h*^2^ from 0 to 1 according to ref ^11^), which reflects 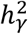 estimates reported in previous African American samples ^30^, and performed admixture mapping to compute summary statistics. Overall, we found HAMSTA produced approximately unbiased estimates of 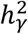 (**Figure 1a**), irrespective of the sparsity of causal markers (**Supplementary Figure 2**). We observed that the summary statistics-based estimates from HAMSTA were highly correlated with those computed from individual-level data using BOLT-REML (**Figure 1b**), suggesting that when stratification bias is not present, there is no loss in accuracy across data settings. Next, to compare our method with existing summary statistics-based methods, we applied LD score regression (LDSC; see **Methods**) and observed LDSC produced biased estimates exhibited large standard errors (**Supplementary Figure 3**). Importantly, we found LDSC estimates remained biased after re-estimating “LAD scores” using a larger window size of 20-cM (**Supplementary Figure 3**). Next, we varied effect of global ancestry while fixing the 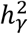 and PVE by ancestral stratification and found HASMTA 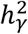 estimates remained unbiased (**Figure 1c**). Together, our results suggest that when stratification does not inflate summary statistics, HAMSTA provides unbiased estimates of 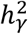, unlike existing summary-based approaches.

**Figure 1.**
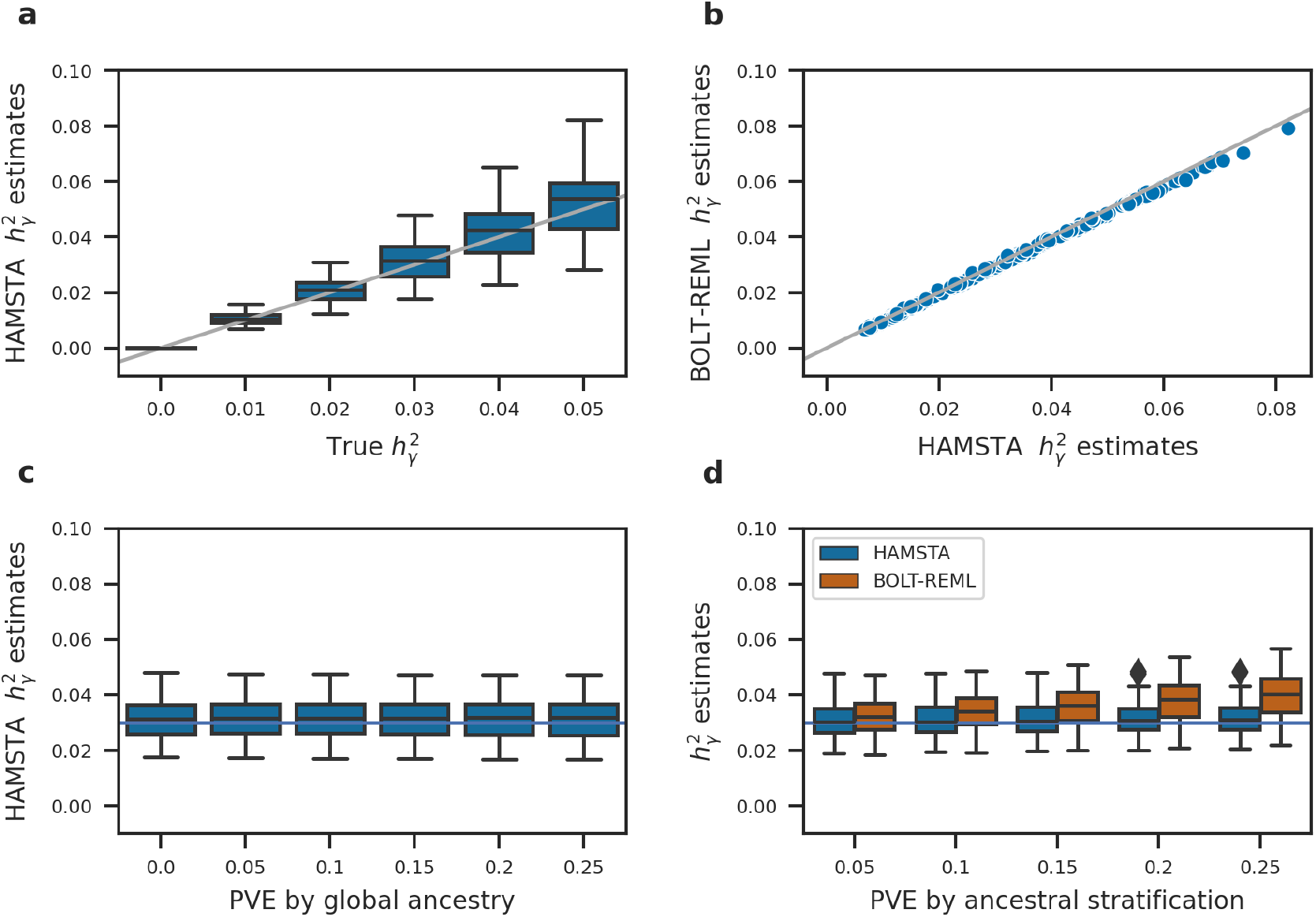
Simulation results from 50,000 admixed individuals and phenotypes under different levels of variance explained by local ancestry, global ancestry and ancestral stratification. The box plots show the range and quartiles of the estimates. a) Results of 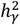 estimation when varying true 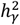. Phenotypic variance explained (PVE) by global ancestry and ancestral stratification were set to 0. A gray identity line is plotted. b) Comparison of 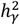 estimates between HAMSTA and BOLT-REML applied to simulation data when true 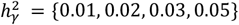 in figure a. c) Results when varying the PVE by global ancestry, setting 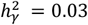 (horizontal line) and PVE by ancestral stratification = 0. d) Comparison of 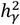 estimates between HAMSTA and BOLT-REML under various levels of ancestral stratification. True 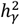 were fixed at 0.03 (horizontal line).

Next, we sought to evaluate HAMSTA in presence of ancestral stratifications. We determined that the 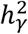 estimates in our method were more robust to the presence of unadjusted ancestral stratification (**Figure 1d**). In contrast, BOLT-REML, where the inference model is not aware of ancestral stratification, produced biased results as the PVE by ancestral stratification increases.

Further, we demonstrate that our method is still robust to other scenarios of structures in the ancestral populations (**Supplementary Figure 4**). We explored the cases where i) both ancestral populations are structured, ii) the proportion of ancestries from the subpopulations are unequal in the admixed population, ii) the subpopulations are introduced to the admixture event at different times. In all the scenarios, the unbiasedness of our estimator is not affected by the ancestral stratification.

Overall, we demonstrated HAMSTA provides unbiased estimates of 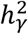 under various levels of effects from local ancestry, global ancestry, and stratification in ancestral populations.

### HAMSTA estimates inflation in admixture mapping statistics due to stratification

Having established the unbiasedness in 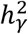 estimates, we next sought to evaluate the ability of HAMSTA to identify inflation in admixture mapping statistics due to ancestral population stratification. Specifically, intercepts estimated by HAMSTA can be tested against the null (i.e., 1) to evaluate stratification bias. Overall, we observed HAMSTA produced estimates greater than 1 as the PVE by ancestral stratification increased (**Figure 2a**), demonstrating the ability of HAMSTA inferred intercepts to capture stratification-induced inflation. Although we noted similar trends in other measures of inflation, including mean *χ*^2^, genomic inflation factor *λ*_*GC*_, their inability to distinguish between polygenicity and confounding prevent their use for complex disease analyses ^13^. Next, we evaluated the ability of LDSC to identify stratification in admixture mapping statistics through its intercept estimates and observed biased results with large variability (**Supplementary Figure 4**). We observed HAMSTA to have significantly greater power to detect stratification bias compared with LDSC (**Figure 2c**). For example, HAMSTA has 80% power when stratification explains 10% of PVE, compared with 5% power of LDSC. These relative differences in performance held when we increased the LAD score window size for LDSC **(Supplementary Figure 4)**. Overall, HAMSTA provides unbiased estimates of inflation in admixture mapping statistics due to ancestral bias and has greater power to reject its null compared to alternative approaches.

**Figure 2.**
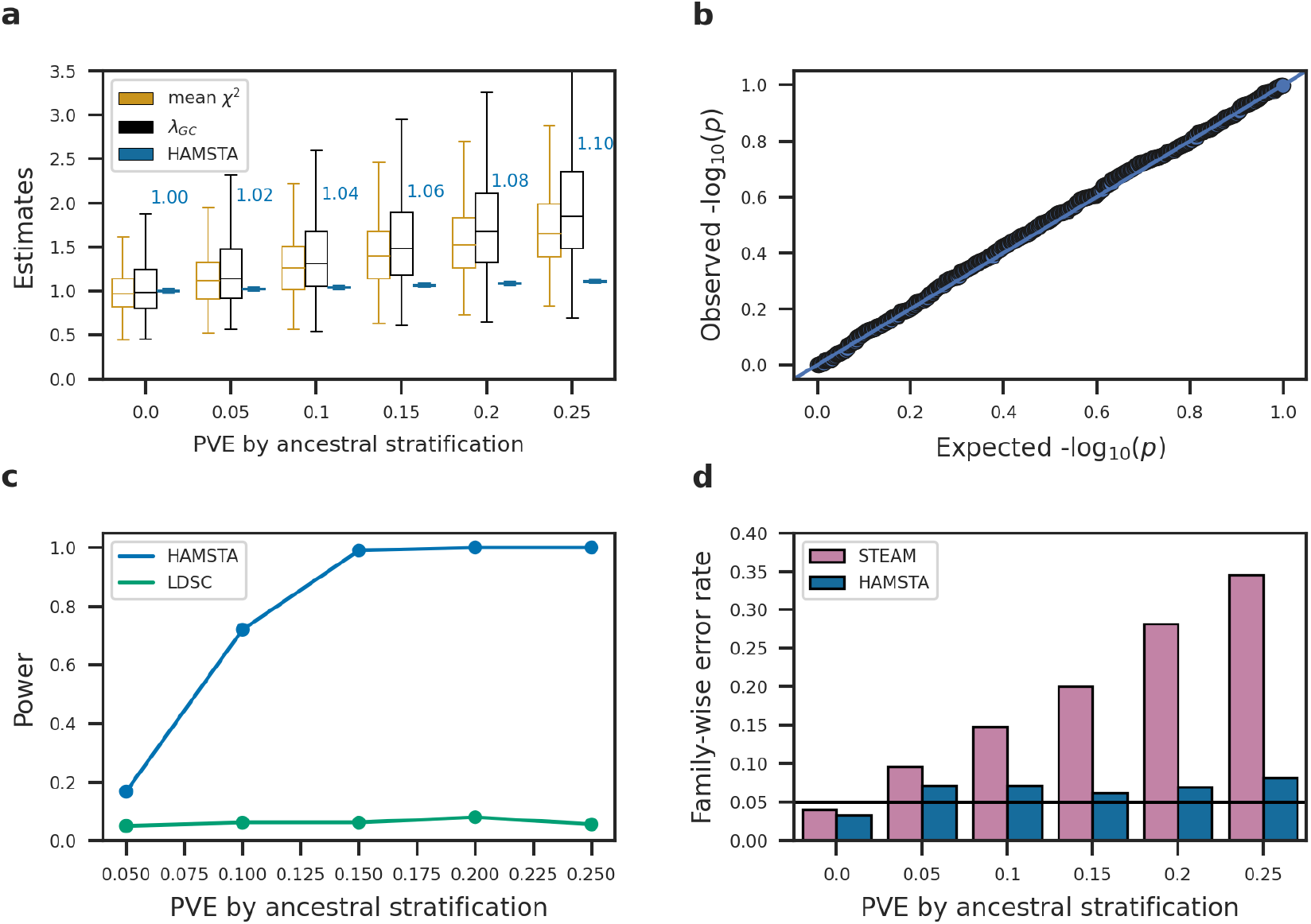
Evaluating ancestral stratification by HAMSTA in 500 simulation replicates. We simulated 50,000 admixed individuals and phenotypes under various levels of variance explained by local ancestral stratification. The true 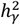 is set to zero. a) Ancestral stratification is reflected by measures of test statistic inflation. The average estimates of HAMSTA’s intercepts are labeled. b) Quantile-quantile plot of test statistics for the test for ancestral stratification. c) Power comparison between HAMSTA and LDSC in detecting ancestral stratification. The p value cutoff for each approach was determined such that the significance level = 0.05 in null simulation d) Family-wise error rate before and after correcting p-value cutoff in admixture mapping using the estimated intercepts.

### HAMSTA improves estimation of p-value thresholds to control family-wise error rate

As the number of approximately independent ancestry blocks depends on the demographic history of the population being studied, there is no universal threshold to determine genome-wide significance in admixture mapping studies. Admixture mapping often relies on permutation-based approaches to estimate the FWER, however these approaches can be computationally intractable for large datasets. Although a recently developed summary-static sampling scheme (STEAM) bypasses the need for individual-level permutations and speeds up the FWER estimation^20^, its assumption that there exists no inflation in the test statistics may be unmet in the presence of population structure and polygenicity.

Here, we demonstrated inferences from HAMSTA can be leveraged to produce significance thresholds for association testing to achieve calibrated family-wise error rates (FWER) compared with STEAM. First, when PVE due to stratification is zero, we found STEAM and HAMSTA estimated similar significance thresholds (HAMSTA mean: 1.12 × 10^−4^; STEAM: 1.57 × 10^−4^), yielding comparable FWER at ∼5% (**Figure 2d**), which suggests that HAMSTA-based FWER estimates do not deflate overall power despite increased model complexity. Importantly, in presence of ancestral stratification, we found HAMSTA estimates resulted in approximately calibrated FWERs unlike STEAM, which produced a considerable number of false positive associations (**Figure 2d, Supplementary Figure 6**). For example, when PVE due to stratification is 0.25, HAMSTA estimates resulted in FWER of 8% compared to the FWER of 34% from STEAM. Together, these findings demonstrate that intercepts estimated by HAMSTA can be incorporated into significance threshold estimation, producing better calibrated FWERs and therefore reducing false positive findings.

### Application to African American in the PAGE study

To illustrate the ability of HAMSTA to estimate 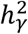 from summary data, we applied it to admixture mapping summary statistics of 20 quantitative phenotypes computed from the African American participants in PAGE study^21^ (mean N = 8383, SD N = 3901; see **Methods**). Briefly, we performed admixture mapping using 101,292 markers adjusting for the study center, global ancestry, and phenotype-specific covariates. The average genomic inflation factor *λ*_*GC*_ across phenotypes is 1.53 (SD = 0.64). Next, we applied HAMSTA to generated summary statistics to infer 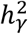 and evaluate potential stratification biases. To estimate *h*^2^ from 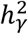, we estimated the average African ancestry to be 78% and *F*_*STC*_ = 0.12 from the admixed individuals in PAGE and reference individuals from HGDP and 1000 Genomes.

We estimated the 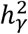 ranges from 0.0025 for systolic blood pressure to 0.033 for height (mean 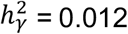; SE = 9.2 × 10^−4^) across the 20 phenotypes, of which 13/20 were individually significantly different from 0 (nominal p-value < 0.05 in **Supplementary Table 1**). Translating 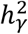 to estimates of *h*^2^, we observed the *h*^2^ ranging from 0.062 for systolic blood pressure to 0.85 for height (mean *h*^2^ = 0.30; SE = 0.023), of which 13/20 were individually significant. We found these results were robust to different values of *F*_*STC*_ (see **Supplementary Table 1)**.

Consistent with the simulation results, HAMSTA estimates were correlated more strongly with BOLT-REML estimates (r = 0.99, **Figure 3**) than those computed from LDSC (r = 0.44) (**Supplementary Figure 7**) This was largely attributable to statistical precision, with standard errors in HAMSTA estimates (range from 0.0023 to 0.014, mean = 0.0058) being slightly greater those from BOLT-REML (range from 0.0021 to 0.0076, mean = 0.0042), and noticeably lower than those computed from LDSC (range from 0.0064 to 0.021, mean = 0.012). Since 5/20 phenotypes had limited sample sizes (N<5,000), which is known to impact the performance of BOLT^26^, we also estimated 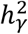 using GCTA. Of the 16 estimates computed by GCTA that converged, we observed they were in general bounded by the estimates by HAMSTA and BOLT-REML (**Supplementary Figure 8**). Overall, we find that HAMSTA estimates of 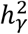 are consistent with those computed from individual-level approaches in real data, while requiring much less computation time for the inference step (49 seconds for HAMSTA versus 51 minutes for GCTA).

**Figure 3.**
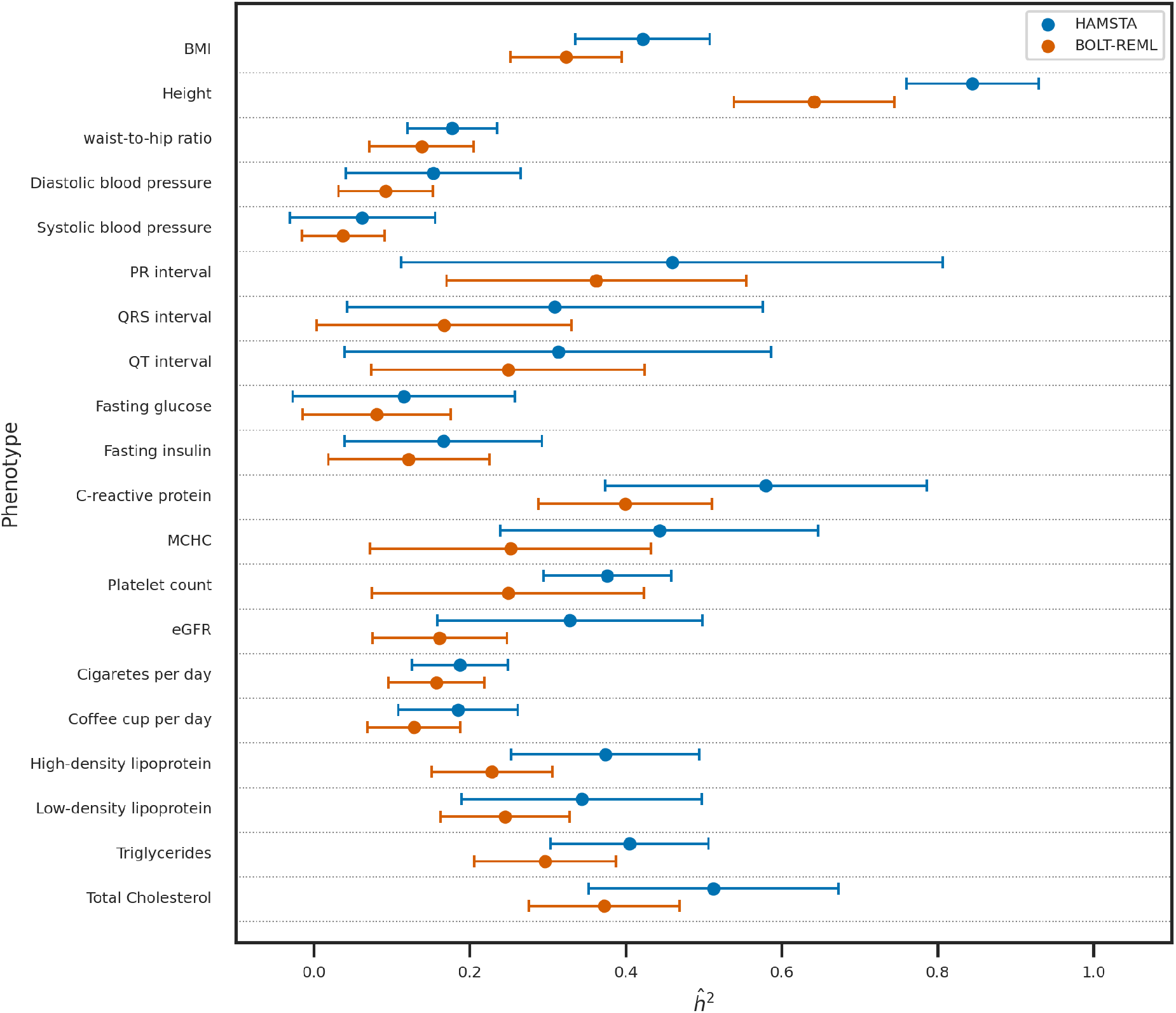
Comparison of 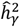 -based 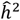 between HAMSTA and BOLT-REML for the 20 quantitative traits in African American in PAGE. Results on 20 PAGE quantitative traits. Comparison between the estimates from HAMSTA, and BOLT-REML. Each point shows the 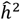, and the lengths of the error bars represent the standard errors.

To substantiate the translated *h*^2^ estimates computed from HAMSTA, we compared with previous *h*^2^ estimates reported from admixed individuals ^11^ as well as those from twin studies. Overall, we found our *h*^2^ estimates are significantly correlated with the previously reported 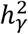 *-*based estimates ^11^ (r = 0.84, p=0.03). Focusing on height, and BMI, HAMSTA estimated 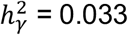 (se: 3.4 × 10^−4^) and 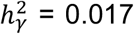 (3.4 × 10^−4^) respectively, corresponding to *h*^2^ of 0.85 (0.085) and 0.42 (0.086) respectively. The estimated height *h*^2^ was similar to the *h*^2^ = 0.68 - 0.84 in twin studies ^31^, whereas the estimated BMI *h*^2^ was smaller than the *h*^2^ = 0.57 - 0.77 in twin studies ^32^ and higher than the *h*^2^ = 0.30 in an estimation from whole-genome sequence data in European ancestry populations^33^.

HAMSTA estimated intercepts suggested limited evidence for inflated summary statistics due to ancestral stratification in the admixture mapping (range from 0.97 to 1.01, average = 0.99; **Supplementary Table 1**), with 0/20 phenotypes differing significantly from the expectation of 1. Although LDSC suggested no significant deviation of intercepts from 1 (range from 0.18 to 1.95, average = 1.07), individual intercepts varied more greatly under LDSC (mean SE = 0.34), than those computed under HAMSTA (mean SE = 5.6 × 10^−3^) (**Supplementary Table 1**).

Since in simulation we demonstrated that the significance threshold for admixture mapping corresponding to FWER of 5% is sensitive to ancestral stratification, we estimated the thresholds based on the HAMSTA intercepts. Under no ancestral stratification (i.e. intercept = 1), HAMSTA estimated the significance threshold required to be 2.80 × 10^−5^, which agrees with the threshold of 2.10 × 10^−5^ reported by STEAM for African American^20^. Based on the estimated intercepts in HAMSTA for the 20 phenotypes, the estimated thresholds range from 2.70 × 10^−5^ to 3.52 × 10^−5^. To conclude, HAMSTA found no evidence of inflation in admixture mapping statistics and provided estimates for 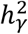 and hence *h*^2^ of the complex traits of African American in PAGE study.

## Discussion

In this study, we demonstrated the use of summary statistics from admixture mapping to quantify the contribution of genetic variations to a trait. We developed a tool, HAMSTA, that unbiasedly estimate 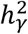 under the various trait architecture, including in the presence of unknown population stratification in ancestral populations. Using the summary statistic-based approach, HAMSTA distinguishes the effect tagged by local ancestry on test statistics from unknown confounding biases. We also demonstrated that the estimated biases could be used to correct the significance threshold such that FWER are better controlled. Lastly, we applied HAMSTA to real-world data, showing that it can recover the 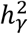 and hence *h*^2^ from admixture mapping summary statistics.

Our method addresses several limitations in existing approaches estimating 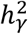. First, because of the long-range correlations between local ancestry markers, LDSC requires a large window size to capture correlations with distant effect markers. Also, regression weights may not be sufficient to solve the problem of correlated *χ*^2^ statistics, which could lead to inefficient estimation ^34^. Our analysis shows that the efficiency can be improved when admixture mapping test statistics are rotated to an independent space. Second, REML could provide an unbiased estimate, but we showed in simulation that it is susceptible to ancestral stratification. Also, it is computationally expensive as the sample size increases. In real data analysis, the REML approach in GCTA failed to converge in waist-to-hip ratio, QT-interval, cigarette-per-day, and HDL. In contrast, we showed that HAMSTA would be a more robust approach to ancestral stratification and has no convergence problem in our analysis. Finally, existing methods assume uniform test statistics inflation although it has been shown that this assumption could be inaccurate ^35,36^. HAMSTA relaxes this assumption by allowing multiple intercepts to represent non-uniform inflation. Overall, HAMSTA offers advantages over existing methods in the above aspects.

We are aware of several limitations of HAMSTA. First, HAMSTA only provides estimates of heritability explained by local ancestries in two-way admixtures, which may limit the use of the method in admixed populations with more than two ancestral populations. Currently, the relationship between 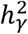 and *h*^2^ are only established in two-way admixed populations such as African American, but models for 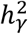 multi-way admixture has not yet been proposed. Incorporating the contribution of multiple ancestries in 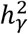 and *h*^2^ will be a possible extension in the future. Second, the standard error of HAMSTA 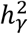 is larger than that from methods that use individual-level data like BOLT-REML (mean SE=0.0058 in HAMSTA versus mean SE=0.0042 in BOLT-REML). Nevertheless, HAMSTA 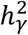 is robust to ancestral stratification, unlike BOLT-REML showing upward biases in the 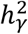 estimates (**Figure 1d**). Third, HAMSTA only models summary statistics computed from linear regression on quantitative traits. The scope of this study is not extended to modeling binary traits. Future work can explore phenotypes under the liability-scale model and evaluate the use of summary statistics from logistic regression models. Lastly, since HAMSTA relies on an accurate LAD, factors that the LAD depends on, such as global ancestries, could potentially impact the accuracy of the estimates. These factors are required to be adjusted for when estimating the LAD.

In summary, our work opens a direction of summary statistics analysis in admixture mapping studies. Our method will facilitate studies of genetic architecture in large cohorts of admixed populations.

## Acknowledgements

This work was funded in part by National Institutes of Health (NIH) under awards R01HG012133 and R35GM142783.

## Web Resources

BOLT-REML, https://alkesgroup.broadinstitute.org/BOLT-LMM/BOLT-LMM_manual.html

GCTA, https://cnsgenomics.com/software/gcta/

GNOMAD HGDP and 1KG subsets, https://gnomad.broadinstitute.org/downloads#v3-hgdp-1kg

LDSC, https://github.com/bulik/ldsc

MSPRIME, https://github.com/tskit-dev/msprime

PLINK, https://www.cog-genomics.org/plink/

RFMIX2, https://github.com/slowkoni/rfmix

STEAM, https://github.com/kegrinde/STEAM

## Data and code availability

The codes for HAMSTA are available at https://github.com/tszfungc/HAMSTA

