## Supplemental Figures for "Estimating heritability explained by local ancestry and evaluating stratification bias in admixture mapping from summary statistics"

### PAGE Phenotype Transformation

The phenotype values were filtered and adjusted according to the original PAGE analysis [1]. For example, After transformation, the phenotypes were then used for admixture mapping with covariates (summarized below).

| Phenotypes | Adjustments | Transformation | Admixture mapping covariates |
|---|---|---|---|
| BMI | Residualized by age, sex age*sex | Inverse normal transformation | Study, global African ancestry |
| Height | Residualized by age, sex age*sex | Inverse normal transformation | Study, global African ancestry |
| Waist-to-hip ratio | Residualized by age, sex age*sex | Inverse normal transformation | Study, global African ancestry |
| Diastolic blood pressure | +10 if use of antihypertensive medication | Winsorization +/- 6 standard deviations | Study, global African ancestry, BMI |
| Systolic blood pressure | +15 if use of antihypertensive medication | Winsorization +/- 6 standard deviations | Study, global African ancestry, BMI |
| PR interval | None | None | Study, global African ancestry, systolic blood pressure, BMI, age, sex, use of beta-adrenergic blocking agents |
| QRS interval | None | None | Study, global African ancestry, systolic blood pressure, BMI, age, sex |
| QT interval | None | None | Study, global African ancestry, heart rate, age, sex |
| Fasting glucose | Residualized by age, sex age*sex, study, smoking status, bmi | Rank normalization | Study, global African ancestry |
| Fasting insulin | Residualized by age, sex age*sex, study, smoking status, bmi | Rank normalization | Study, global African ancestry |
| C-reactive protein (CRP) | None | natural logarithm of one plus | Study, global African ancestry, age at measurements, smoking status, sex |
| MCHC | None | None | Study, global African ancestry, age at measurements, smoking status, sex |
| Platelet count | None | None | Study, global African ancestry, age at measurements, smoking status, sex |
| eGFR | None | No transformation | Study, global African ancestry, age, sex |
| Cigarette per day | None | natural logarithm of one plus | Study, global African ancestry, age, sex |
| Coffee cup per day | None | natural logarithm of one plus | Study, global African ancestry, age, sex |
| HDL | Largest adjustment from:<br>statins: -2.3;<br>fibrates: -5.9;<br>bile acid sequestrants: -1.9;<br>niacin: -9.9,<br>cholesterol absorption inhibitors: 0 | None | Study, global African ancestry, age at lipid measurement, sex |
| LDL | Largest adjustment from:<br>statins: +49.9;<br>fibrates: +40.1;<br>bile acid sequestrants: +40.5;<br>niacin: +24.7,<br>cholesterol absorption inhibitors: +40.5 | None | Study, global African ancestry, age at lipid measurement, sex |
| Triglyceride | Largest adjustment from:<br>statins: +18.4;<br>fibrates: +57.1;<br>bile acid sequestrants: 0;<br>niacin: +89.4,<br>cholesterol absorption inhibitors: 0 | None | Study, global African ancestry, age at lipid measurement, sex |
| Total Cholesterol | Largest adjustment from:<br>statins: +52.1;<br>fibrates: +46.1;<br>bile acid sequestrants: 0;<br>niacin: +34.6,<br>cholesterol absorption inhibitors: +40.5 | None | Study, global African ancestry, age at lipid measurement, sex |

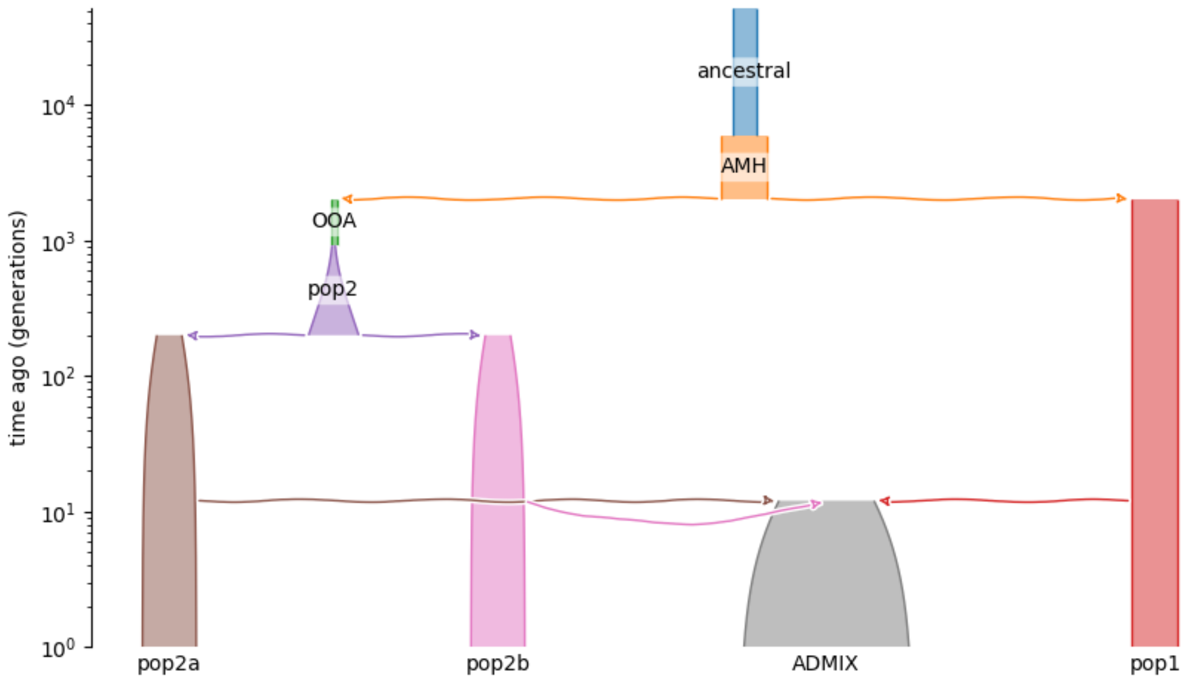


**Supplementary Figure 1**
The demography in the simulation study. Timeline is shown on the y-axis in log scale. Arrows represent the direction of migrations of between populations. Ancestral: Ancestral equilibrium. AMH: Anatomically modern humans. OOA: Out-of-Africa. ADMIX: Admixed.

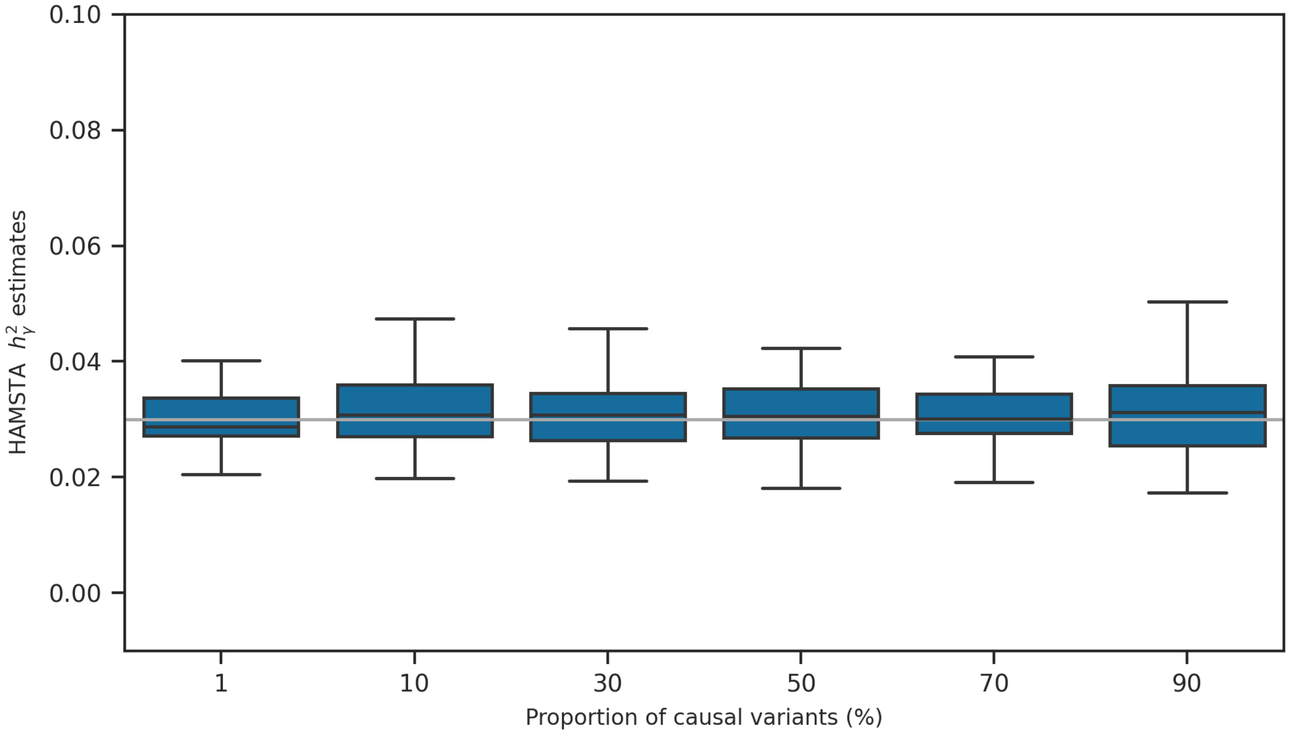


**Supplementary Figure 2**

Boxplots showing HAMSTA $h_\gamma^2$ estimates when varying the number of causal variants out of the complete 20,000 markers. The boxplot consists of 50 simulation replicates. The true $h_\gamma^2$ is set at 0.03 shown in the gray horizontal line.

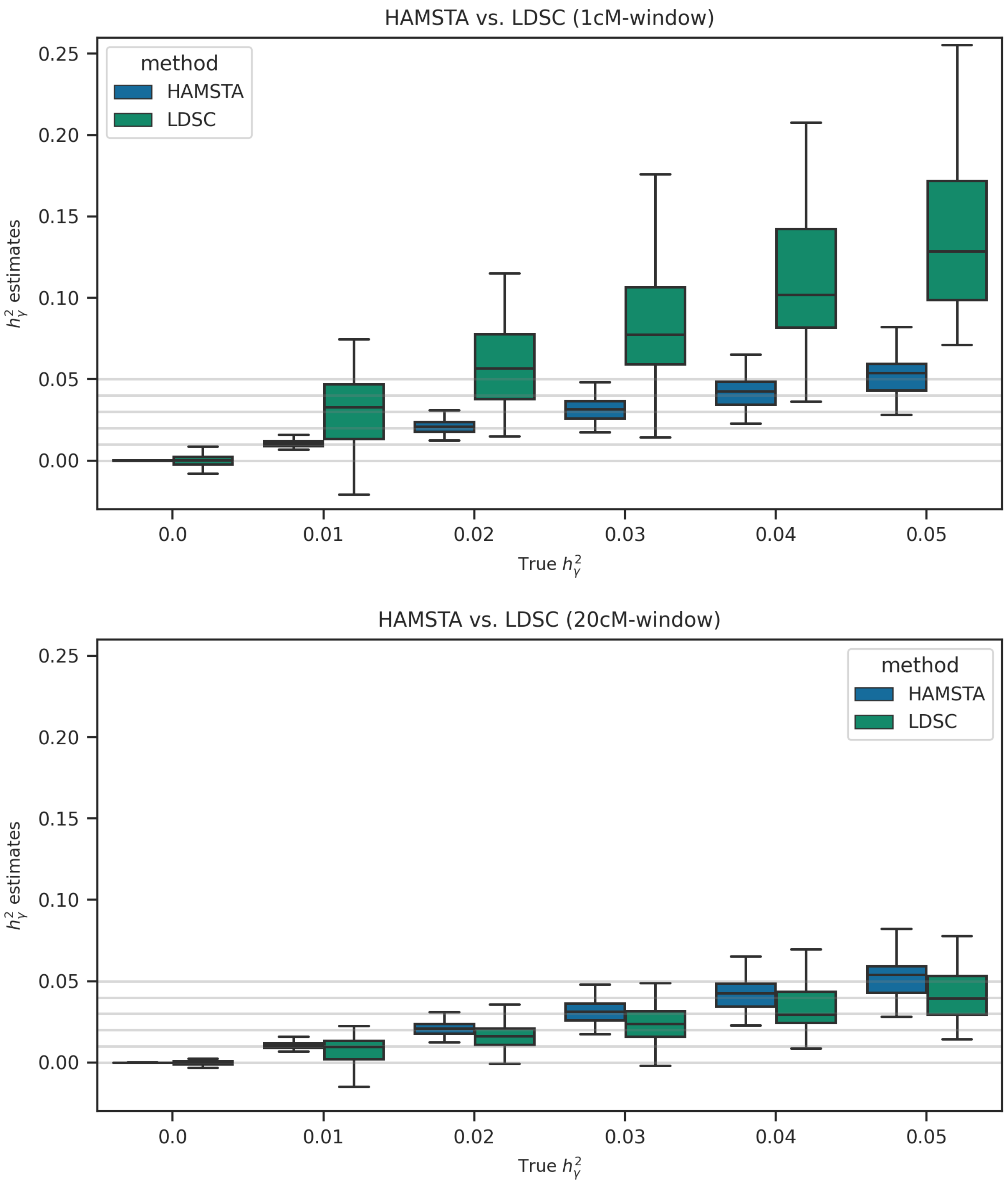


**Supplementary Figure 3**

Boxplots comparing $h_\gamma^2$ estimates from HAMSTA and LDSC at various true $h_\gamma^2$. Each $h_\gamma^2$ consists of 50 simulation replicates. The true $h_\gamma^2$ are set at {0.0, 0.01, 0.02, 0.03, 0.04, 0.05} shown in the gray horizontal lines. In the upper and lower plots, LAD scores are computed in 1cM- and 20cM-window respectively.

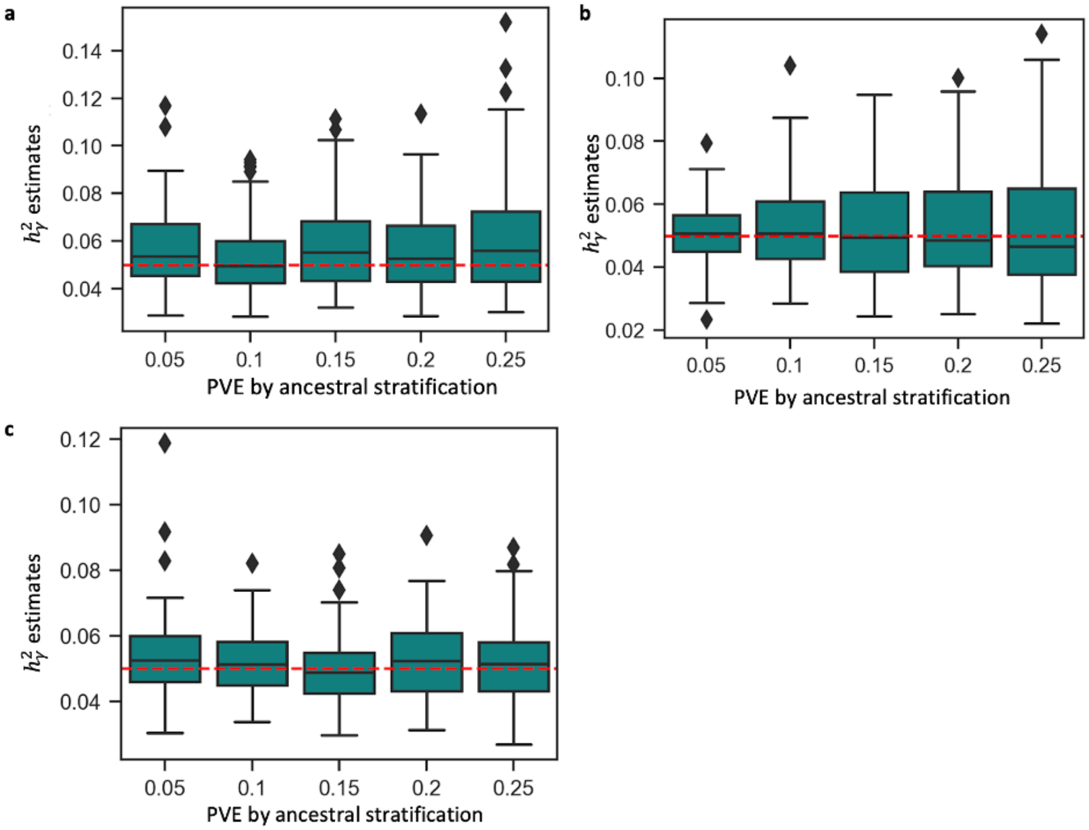


**Supplementary Figure 4**

Boxplots showing HAMSTA $h_\gamma^2$ estimates under various levels of variance explained by local ancestral stratification in 50 replicates under other demographic models in which a) population structure is present in both ancestral populations, b) subpopulations in the structured ancestral population contributed 4% and 16% ancestral proportion, and c) subpopulations in the structured ancestral population migrated to the admixed population separately 12 and 8 generations ago. The true $h_\gamma^2$ is set at 0.05 shown in the red horizontal line. Points that deviate from upper or lower quartiles by 1.5 times of interquartile range are marked as diamonds.

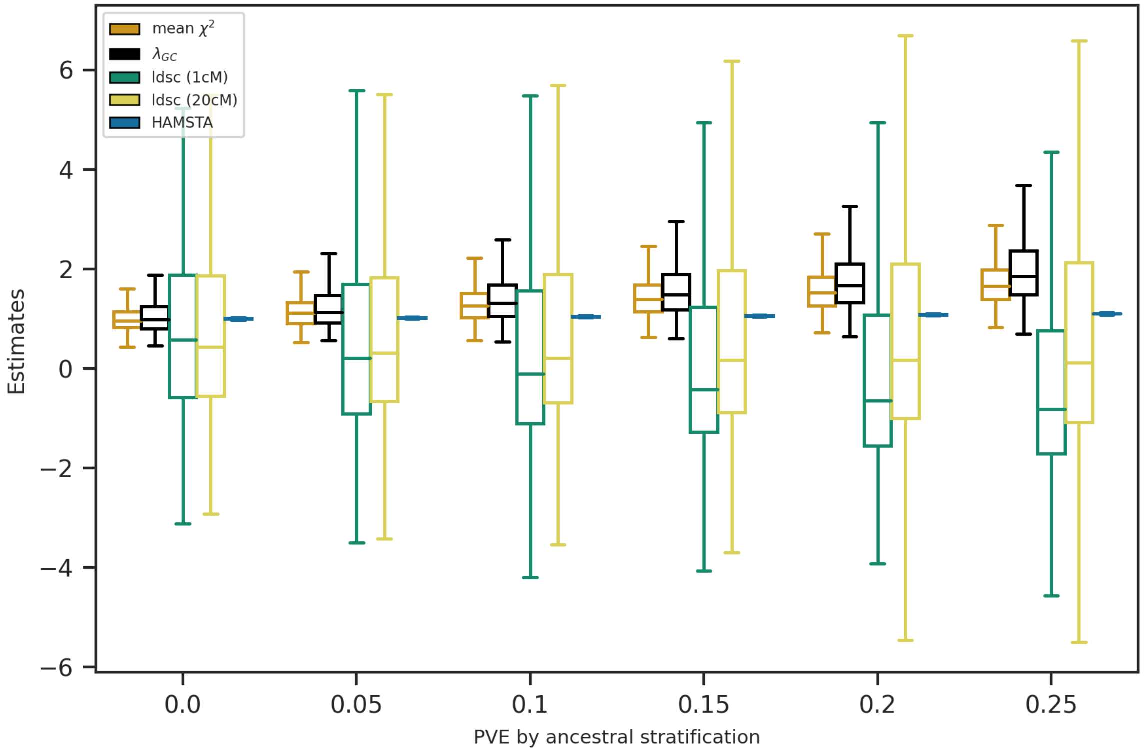


**Supplementary Figure 5**

Boxplots comparing different measures of test statistics inflation at various levels of PVE by ancestral stratification. The LAD scores in LDSC were computed with window sizes of 1cM and 20cM. The mean intercepts from HAMSTA were used to represent the inflation for one simulation.

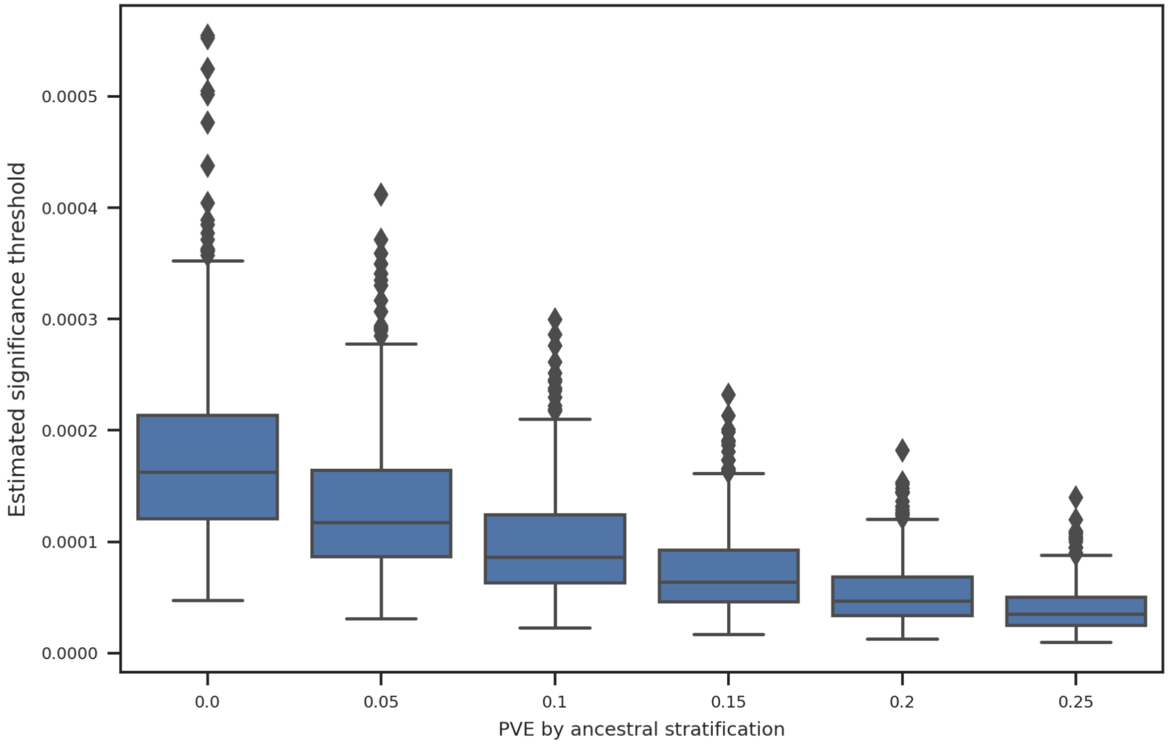


**Supplementary Figure 6**

Estimated significance threshold for admixture mapping at various levels of PVE by ancestral stratification in 500 simulations with $h_\gamma^2 = 0$. Points that deviate from upper or lower quartiles by 1.5 times of interquartile range are marked as diamonds.

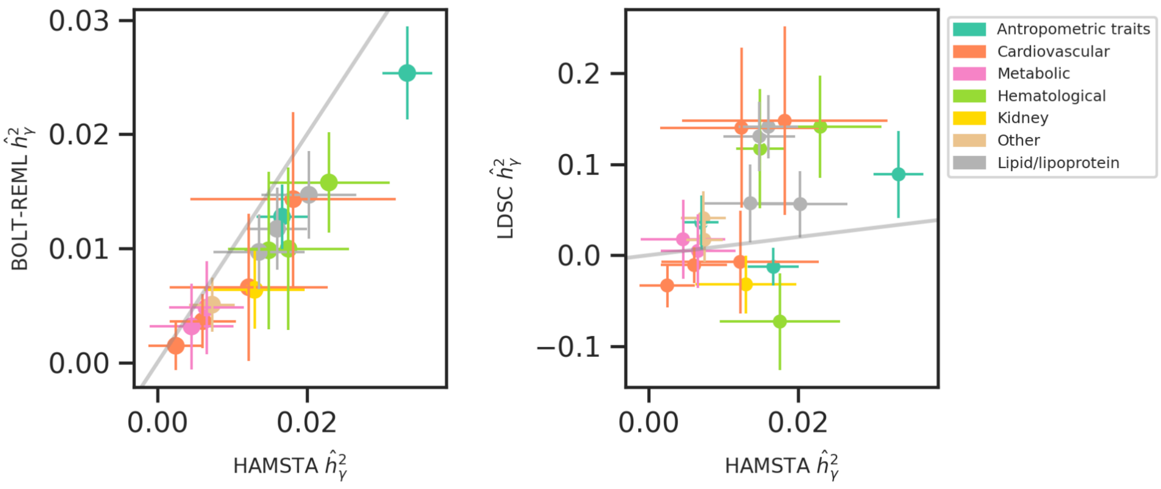


**Supplementary Figure 7**

Results on 20 PAGE quantitative traits. Comparison of the estimates between HAMSTA and BOLT-REML, and those between HAMSTA and LDSC. The points show the point estimates of $h^2$. The lengths of the error bars represent the standard errors.

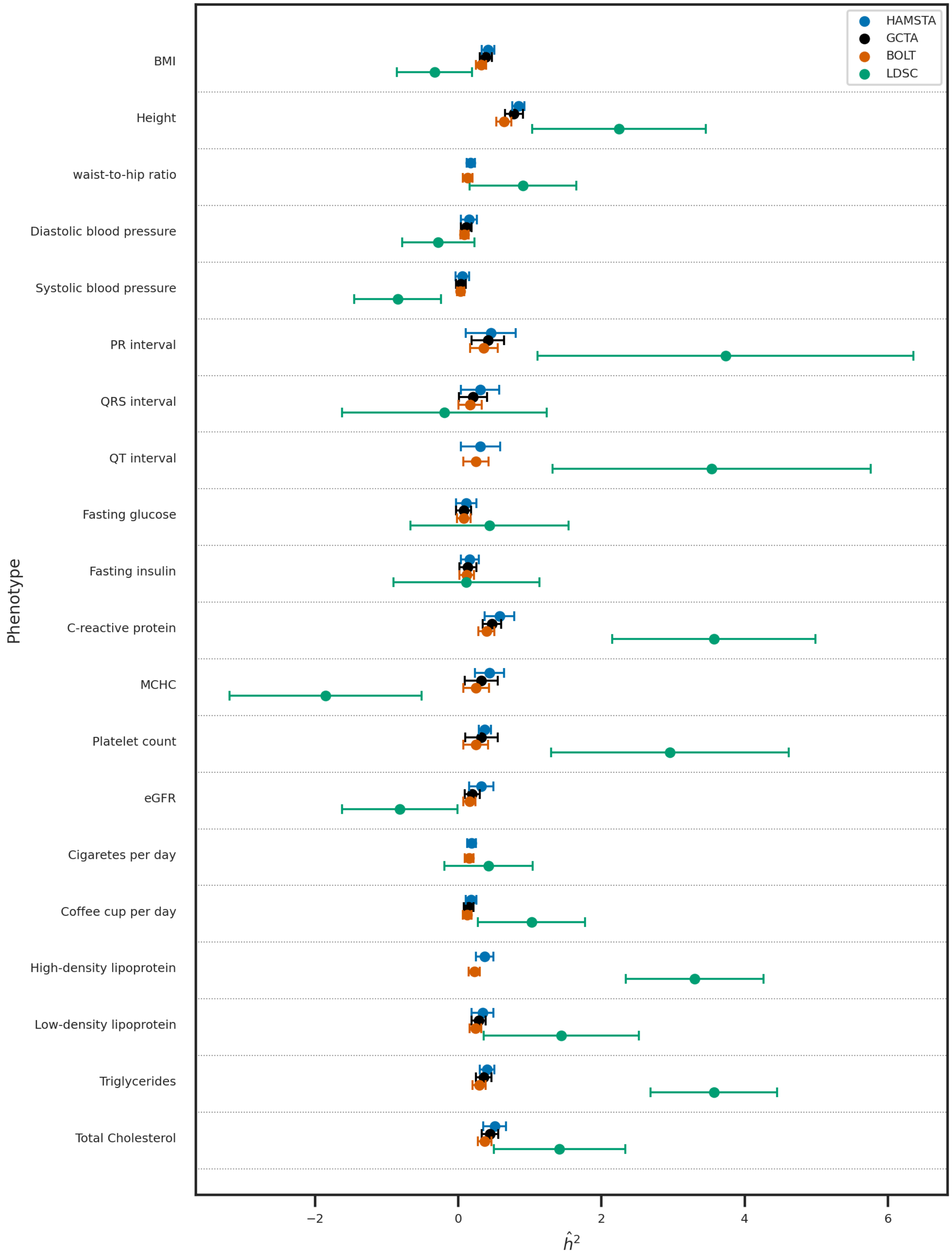


**Supplementary Figure 8**

Results on 20 PAGE quantitative traits. Comparison between the estimates from HAMSTA, BOLT, and GCTA, and LDSC. The points show the point estimates of $h^2$. The lengths of the error bars represent the standard errors.
